# Revealing the unique features of each individual’s muscle activation signatures

**DOI:** 10.1101/2020.07.23.217034

**Authors:** Jeroen Aeles, Fabian Horst, Sebastian Lapuschkin, Lilian Lacourpaille, François Hug

## Abstract

There is growing evidence that each individual has unique movement patterns, or signatures. The exact origin of these movement signatures however, remains unknown. We developed an approach that can identify individual muscle activation signatures during two locomotor tasks (walking and pedalling). A linear Support Vector Machine was used to classify 78 participants based on their electromyographic (EMG) patterns measured on eight lower limb muscles. To provide insight into decision making by the machine learning classification model, a Layer-wise Relevance Propagation (LRP) approach was implemented. This enabled the model predictions to be decomposed into relevance scores for each individual input value. In other words, it provided information regarding which features of the time-varying EMG profiles were unique to each individual. Through extensive testing, we have shown that the LRP results, and by extent the activation signatures, are highly consistent between conditions and across days. In addition, they are minimally influenced by the dataset used to train the model. Additionally, we proposed a method for visualising each individual’s muscle activation signature, which has several potential clinical and scientific applications. This is the first study to provide conclusive evidence of the existence of individual muscle activation signatures.

## Introduction

Evolution has resulted in immense variation and specialisation within and between the phyla of the kingdom Animalia (1). For example, humans have evolved distinct movement patterns (e.g., gait) that separate us from non-human primates (2). However, even within the human species, different movement patterns are often observed when individuals execute the same motor task (3). Recent studies have taken advantage of machine learning techniques to show that computer models can identify individuals based on kinematic or kinetic data generated during walking (4–6). This suggests that the participants exhibited discernible differences in their movements and, therefore, that patterns of movement not only vary between individuals but are unique to each individual. This has led to the concept of individual movement signatures, defined as distinctive patterns or characteristics by which an individual can be identified (7, 6). Identifying the origin of such movement signatures is crucial for understanding the control of movement in health and disease (8).

As movement is the result of the coordinated activation of multiple muscles, individual movement signatures may originate from unique muscle activation patterns. However, given the complex interplay between biomechanical properties of contracting muscle, tendon, and the skeletal system, it is also possible that different movement patterns are achieved with similar muscle activation strategies among individuals. Yet, the existence of individual muscle activation signatures was recently suggested by Hug et al. (9) who demonstrated that a machine learning algorithm can accurately identify individuals based on the activation profiles of eight muscles measured with surface electromyography (EMG) during walking and pedalling. However, despite the power of machine learning approaches for identification of individuals, their main limitation is that they do not provide any information regarding which features are important to the decision-making process of the model (10, 11). In other words, it remains unknown which features of muscle activation patterns are used to identify individuals and therefore what makes each individual unique.

The limitation of classical approaches can be overcome by techniques that provide insight into the decision-making process of the machine learning classification model, such as Layer-wise Relevance Propagation (LRP) (10). The implementation of this technique in the field of computer vision has made it possible to highlight which pixels are crucial in the classification of images and allows us to understand and interpret the decision-making process of the models (12). Developing this approach for identifying muscle activation signatures is critical, as this will allow us to unravel the physiological origin and relevance of individual signatures.

In this study, we extended the analyses conducted by Hug et al. (9) by using the LRP technique to decompose the predictions of a Support Vector Machine (SVM) classification model based on EMG data collected during pedalling and walking. We considered these two motor tasks for their different mechanical constraints: pedalling, for which the movement velocity, foot trajectory, and torque can be matched between participants; and gait, which is a more natural and less-constrained form of locomotion. The overall aim of this study was to identify the unique features in individual muscle activation signatures. Specifically, the first aim was to determine whether the identification of individuals based on their muscle activation patterns relied on periods when the muscles were most active. The second aim was to test the within-session, between-condition, and between-days reliability of the LRP results and to test the dependency of the results on the dataset used to train the model. We also propose a method for graphical depiction of individual muscle signatures that could be used to easily compare between individuals.

## Materials and Methods

### Experimental Design

The present study used an innovative approach to analyse a dataset that was publicly released by our group (doi.org/10.6084/m9.figshare.8273774) and that was partially analysed in a previous research article *(9).*

Eighty physically active volunteers participated in this study (62 men and 18 women; mean ±standard deviation; age: 23.6 ± 5.4 yrs, height: 176.7 ± 7.8 cm, body mass: 71.3 ± 10.0 kg). Participants had no history of lower leg pain within the previous two months. The institutional research ethics committee, CPP Ile de France XI, approved this study (no. 2018-A00110–55/18020), and all procedures adhered to the Declaration of Helsinki. All participants provided written informed consent.

The experimental protocol has previously been described in detail *(9).* Data were collected during two separate sessions. In the first session (referred to as Day 1), all 80 participants were tested. In the second session (referred to as Day 2), which was held (mean ± standard deviation); 13 ± 10 days after Day 1, 53 participants (12 women and 41 men) were tested again. The session on Day 1 consisted of a series of locomotor tasks: two all-out isokinetic pedalling sprints were used to standardize the intensity of the submaximal pedalling tasks, pedalling at four submaximal intensities, and walking on a treadmill at 1.11 m/s. The order of the tasks (pedalling and walking) and the intensities of the pedalling tasks were randomized for each session. On Day 2, only the submaximal pedalling tasks and the walking task were repeated.

The pedalling task was performed on an electronically braked cycloergometer (Excalibur Sport; Lode, Groningen, the Netherlands) equipped with clipless pedals and cycling shoes. Saddle height, saddle setback, and handlebar height were adjusted to the anthropometry of the participant in order to limit the impact of the pedalling position on the variability of the activation strategies between the participants and between the two test sessions (for more details, see *9*). After familiarizing themselves with the cycloergometer and a standardized warm-up, participants performed two 5-s all-out isokinetic pedalling sprints at 80 rpm, separated by 2 min of rest. The average of the two cycles with the highest power output was considered to be the maximal power output (Pmax). Next, participants pedalled at four different submaximal intensities (80 W, 150 W, 10% of Pmax, and 15% of Pmax; in a randomized order), each at 80 rpm for 90 s, separated by 30-s of rest. Feedback from the target pedalling rate was displayed on a monitor placed in front of the participants. The 150 W condition from Day 1 was selected as the condition for pedalling on which the classification model was trained, and is referred to as the baseline data for pedalling. Data for the 80 W were not needed to address the aims of the present study and therefore are not reported. Participants walked barefoot on a treadmill (Power 795i, Pro-form, France) and first familiarized themselves with the treadmill before starting the experimental task, which consisted of walking at 1.11 m/s for 90-120 s. The Day 1 walking data were selected as the condition for walking on which the classification model was trained and is referred to as the baseline data for walking.

### EMG Recordings

EMG signals were collected from eight muscles on the right leg: vastus lateralis (VL), rectus femoris (RF), vastus medialis (VM), gastrocnemius lateralis (GL), gastrocnemius medialis (GM), soleus (SOL), tibialis anterior (TA), and biceps femoris-long head (BF). For each muscle, a wireless surface electrode (Trigno Flex, Delsys, Boston, USA) was attached to the skin at the site recommended by SENIAM (Hermens et al., 2000). Before attaching the electrodes, the skin was shaved and then cleaned with an abrasive pad and alcohol. Electrodes were secured to the skin with double-sided tape and a tubular elastic bandage (tg®fix, Lohmann & Rauscher International, GmbH & Co. KG, Germany). EMG signals were band-pass filtered (10-850 Hz) and digitized at a sampling rate of 2,000 Hz using an EMG acquisition system (Trigno, Delsys, Boston, USA). A trigger signal indicating either the top-dead centre of the right pedal (pedalling, using a hall effect sensor) or the onset of foot contact (walking, using a force sensing resistor) was recorded on the EMG acquisition system.

### Data Pre-processing

All EMG data were pre-processed offline using Matlab R2015b (MathWorks, USA). Raw EMG signals were first band-pass filtered (20–700 Hz) and were then visually inspected for noise or artefacts. At this stage, some data were discarded due to movement artefacts. In this study, we considered only the participants for whom data were available for both pedalling and walking on Day 1, which left data for 78 participants for analysis.

For both tasks, the first 20 cycles were excluded from the analysis. Then, the first 30 consecutive cycles that were free of any artefacts were selected. The EMG signals were full-wave rectified and low-pass filtered at either 12 Hz (pedalling) or 9 Hz (walking). As explained in detail in Hug et al. (*9*), these cut-off frequencies were selected because they provided the best classification accuracy and fell within the range classically used for smoothing EMG signals during locomotor tasks. For every muscle, data for each cycle were interpolated to 200 data points and normalized to that muscle’s maximal EMG amplitude that was measured within that cycle. This normalization procedure reduced the chance that the classification process was biased toward the muscles that exhibited the highest EMG amplitude. As such, all muscles were equally weighted in the classification process.

### Data Classification

Classification was conducted using Python 3.7 (Python Software Foundation, USA; codes are available at https://github.com/sebastian-lapuschkin/interpretable-emg-signatures). To test the uniqueness of muscle activation strategies during pedalling and walking, we used a concatenated vector of the time-varying EMG profiles of all eight muscles from the baseline data (150 W, Day 1 for pedalling or Day 1 for walking). The ability to distinguish between the EMG profile of one participant versus the EMG profiles of all other participants was investigated in a multi-class classification (participant-classification) setting. First, the classification performance of three machine learning approaches (i.e. (linear) SVM, Multi-layer Perceptron (MLP), and Convolutional Neural Network (CNN)) was compared. As the classification performance of the SVM model was systematically superior and because its computational time was substantially shorter than for MLP and CNN, only the data analysed with the SVM model are reported. The SVM models were trained using a standard quadratic optimization algorithm, with an error penalty parameter of *C* = 0.01 and l2-constrained regularization of the learned weight vector. For the evaluation of classification performance, the prediction accuracy was reported over a stratified 10-fold cross-validation configuration, where in each repetition of the evaluation, eight partitions of the data were used for training (24 cycles), one partition was used for validation (3 cycles), and the remaining partition was used for testing (3 cycles). The 30 cycles of each participant were randomly shuffled and evenly distributed across the partitions such that each unique group of three out of the 30 cycles was included once in the test partition.

### Classification Explanation

Determining which features of the EMG profiles were unique to each individual participant represents the originality of the present work. Specifically, we determined which muscle(s) and at which time period within the pedalling or walking cycles were relevant to the classification of each participant according to the model. To this end, we used the Layer-wise Relevance Propagation (LRP) method *(10)*. LRP decomposes the prediction *f_(x)_* of a learned function *f* given an input vector *x* into time- and muscle-resolved input relevance scores for each discrete input unit. This enables us to explain the prediction of a machine learning model as partial contributions of an individual input value. In other words, LRP indicates which information was used by the model for its prediction of either in favour of (positive input relevance scores) or against (negative input relevance scores) an output class, i.e. participant. This enables the input relevance scores and their dynamics to be interpreted as representation of a certain class, i.e. an individual muscle activation strategy.

A derivation of the LRP Toolbox for Python (version 1.3.0) (13) was used to obtain relevance attributions. Since the models investigated in this study are comparatively shallow and are largely unaffected by detrimental effects such as gradient shattering, we performed relevance decompositions according to LRP-ε, with *ε* = 10 - 5 in all layers. For each cycle, we obtained input relevance scores in its function as a test sample during the crossvalidation procedure. We thus obtained an understanding of how the model generalizes toward each participant based on data that were unseen during training. The relevance scores were exported to Matlab R2019B (MathWorks, USA) in which all further analyses were performed. Pedalling and walking data were analysed separately using the same methodological approach.

Figure 1 shows a graphical overview of the data analysis steps. Unless otherwise stated, the relevance pattern of a single cycle contained all the relevance scores for the 200 time points and for all eight muscles as a single concatenated vector. An example of such a cycle is presented in Fig. 1A for a representative participant. Because negative input relevance scores are difficult to interpret in multi-class classifications, we considered only the positive relevance scores, which identify the features of the input that describe the true participant label exclusively. After excluding negative relevance scores (by replacing them with zeroes; Fig. 1B), the positive relevance scores were normalized to the maximal relevance score achieved for that cycle. This resulted in relevance scores for each of the 30-cycles per participant ranging between 0 and 1 (Fig. 1C). From these 30 cycles, the mean relevance score per feature (i.e. data point) was calculated, which resulted in a mean relevance curve for each participant. The mean relative EMG amplitude was calculated similarly for each participant based on the relative EMG amplitudes of all 30 cycles. For some of the additional analyses, each of the 30 individual cycles were used while for others the mean relevance data per participant were used, which is explicitly specified where applicable. To determine whether high positive relevance occurred during periods of high relative EMG amplitudes (aim 1), we tested the relationship between the relevance score and the relative EMG amplitude. To this end, we calculated the Pearson correlation coefficient on the pooled data of all participants, consisting of the mean relevance data and the mean EMG data for each of the 200 time points and for each participant.

**Fig. 1.**
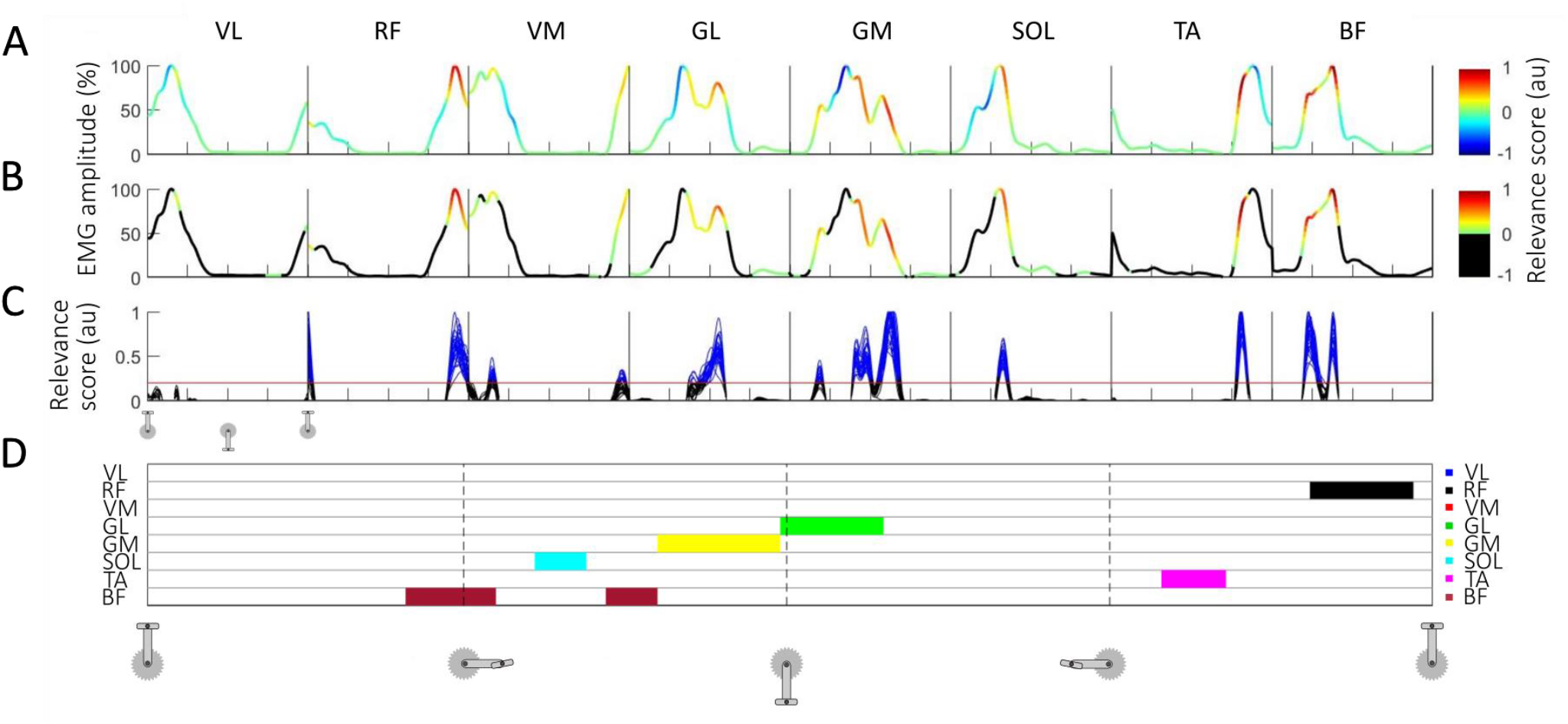
Methodological overview and example results. Graphical overview of the post-Layer-wise Relevance Propagation analyses for an example participant. Data for pedalling (Day 1) start with the pedal on the top dead centre. The x-axis ticks on the top three panels (A-C) and the vertical dashed lines on panel D denote 25, 50, and 75% of the pedalling cycle. Each box on the top three panels (A-C) shows the data from the same cycle(s) but for each muscle. The line in panel A shows the normalised EMG patterns for all eight muscles from a single cycle during pedalling, with the colour scales representing the relevance score from the LRP analysis. Negative relevance scores were discarded by replacing them with zeroes (panel B). In Panel C, the arbitrary threshold of 0.2 is shown as a red horizontal line and all the data above this threshold are coloured blue. Only the data points for which the relevance scores exceeded this threshold in all 30 cycles for a specific timepoint were used to create the signature map shown in panel D (discussed in the methods section Graphical Depiction of the Individual Muscle Activation Signatures). Here, each horizontal line corresponds to a muscle and each muscle is given a colour code that is shown on the right side of panel D (VL = vastus lateralis; RF = rectus femoris; VM = vastus medialis; GL = gastrocnemius lateralis; GM = gastrocnemius medialis; SOL = soleus; TA = tibialis anterior; BF = biceps femoris). For example, several cycles for the VM muscle correspond to data points that exceed the threshold (in blue; panel C) but it can be seen that this did not consistently happen in all 30 cycles and thus we did not consider it to be part of the participant’s consistent muscle activation signature.

### Robustness of the Relevance Scores

For the robustness analysis (aim 2), some participants were excluded due to artefacts in the EMG data. As a result, the data corresponding to 77 participants for pedalling at 10% of Pmax (Day 1) and data corresponding to 49 and 50 participants for pedalling and walking on Day 2, respectively, were used in testing. First, we assessed the reliability of the relevance scores across the 30 cycles (within-session reliability), between pedalling conditions (between-condition reliability), and between Day 1 and 2 (between-day reliability). The baseline data from pedalling at 150 W (Day 1) and walking at 1.11 m/s (Day 1) were used to train and validate the models. Next, we tested the generalization performance of these trained models for participant classification based on the participant-matched test data from pedalling at 10% of Pmax and 15% of Pmax on Day 1 (between-condition reliability) and from pedalling at 150 W on Day 2 and walking on Day 2 (between-day reliability). To assess reliability, the RMSE and Pearson correlation coefficients between the relevance scores were calculated. For the within-session reliability, the relevance scores of each of the 30 cycles were compared with each other (870 combinations per participant) and the average RMSE was determined by calculating the mean of all resulting RMSE values for that participant. The correlation analysis was performed on the same number of combinations. To test the between-condition (pedalling) and between-day (pedalling and walking) reliability, we performed the same analyses as described above but instead used the relevance scores averaged over the 30 cycles for each participant.

Second, we tested the robustness of the relevance scores with respect to the dataset used to train the models in terms of the number of participants included in training and testing the models. To this end, we used the baseline data from pedalling at 150 W on Day 1 and walking at 1.11 m/s on Day 1. We created subsamples with different numbers of participants (steps of 4, from N = 6 to N = 74), resulting in 18 subsample sizes. To avoid a bias based on which of the 78 participants were included, we ran multiple iterations of different combinations of participants. To this end, 10 different combinations of participants were made with only one participant included in all 10 combinations (tested participant) and the other participants selected randomly each time. This was then repeated for each of the 78 participants such that each of them was once the tested participant. For each of these model iterations, we used the above-described ten-fold cross-validation procedure and processed the relevance results from the LRP using the same methods as described above. Each of these iterations produced the relevance scores for each of the 30 pedalling or walking cycles per participant. For further analyses, the mean relevance scores per feature over the 30 cycles were calculated for each participant. Because the results from the different combinations of participants were all highly similar, we further used the mean correlation coefficients of the 10 iterations for each participant and each subsample. Then, the Pearson correlation coefficient between this mean relevance score per participant and that participant’s baseline (N= 78; Day 1) mean relevance score was calculated for each subsample size. In other words, this procedure allowed us to compare the time-varying relevance patterns from the baseline data of each participant to their time-varying relevance patterns identified from different models with a different subset of participants. This step was crucial to discuss the generalization of our results.

### Graphical Depiction of the Individual Muscle Activation Signatures

Even though the relevance scores were normalised to their maximum values during a single cycle, forming a correct interpretation on the magnitude of the relevance scores is not straightforward. For this reason, we decided to only consider the values that were above an arbitrary threshold of 0.2 (out of 1) at each time point in all 30 cycles as part of the signature (Fig. 1C in blue). Even though relevance scores below this threshold were occasionally observed (Fig. 1C), they did not consistently occur for all the cycles, casting doubt on their physiological relevance. Furthermore, features with such low relevance scores (below the threshold), only contribute little to the classification of the participant and it therefore has little impact on the further results. To identify which muscles were the most relevant to identify participants, we determined, for each muscle, the number of participants for whom the mean relevance score exceeded the threshold of 0.2. Further, to provide a graphical depiction of the individual muscle activation signatures, we extracted the periods of the cycle and the muscles for which the relevance score reached the threshold in each of the participant’s 30 cycles (Fig. 1D).

### Statistical Analysis

Mean RMSE and Pearson correlation coefficients were used to assess the robustness of the technique. For each analysis, Pearson correlation coefficients were transformed to Fisher’s Z values, from which the mean and 95% confidence intervals were calculated before transforming the result back to obtain a single mean correlation coefficient and 95% confidence intervals. Statistical significance for all correlations was set at *P < 0.05.*

## Results

### Classification Accuracy

We previously developed an innovative approach for identifying muscle activation signatures based on existing data (*9*). In Hug et al. (*9*), we showed that a classification model is capable of assigning EMG patterns that were measured during pedalling and walking to the corresponding individual with an accuracy up to 99.28%. This result suggested the existence of individual muscle activation signatures. In the current study, we used a similar SVM model, which logically achieved classification accuracies that were extremely close to those reported in Hug et al. (*9*); i.e. (mean ± standard deviation), 99.3 ± 0.5% for pedalling and 98.9 ± 0.5% for walking (results are from the ten-fold cross validation).

The disadvantage of using SVM and other machine learning techniques for classification is that the models provide little information regarding how they arrived at the prediction for individual samples (e.g., individual pedalling or walking cycles). In this work, we used a novel approach based on the LRP technique *(10*) to explain the uniqueness of the time-varying EMG patterns (Fig. 1). LRP enables the decomposition of the predictions made by a machine learning model into relevance scores for each individual input value; i.e. normalised EMG amplitude value. That is, LRP indicates which input values the machine learning model based its prediction on for each participant. In other words, LRP allows us to identify the features of the time-varying EMG profiles that make each individual unique (Fig. 2).

**Fig. 2.**
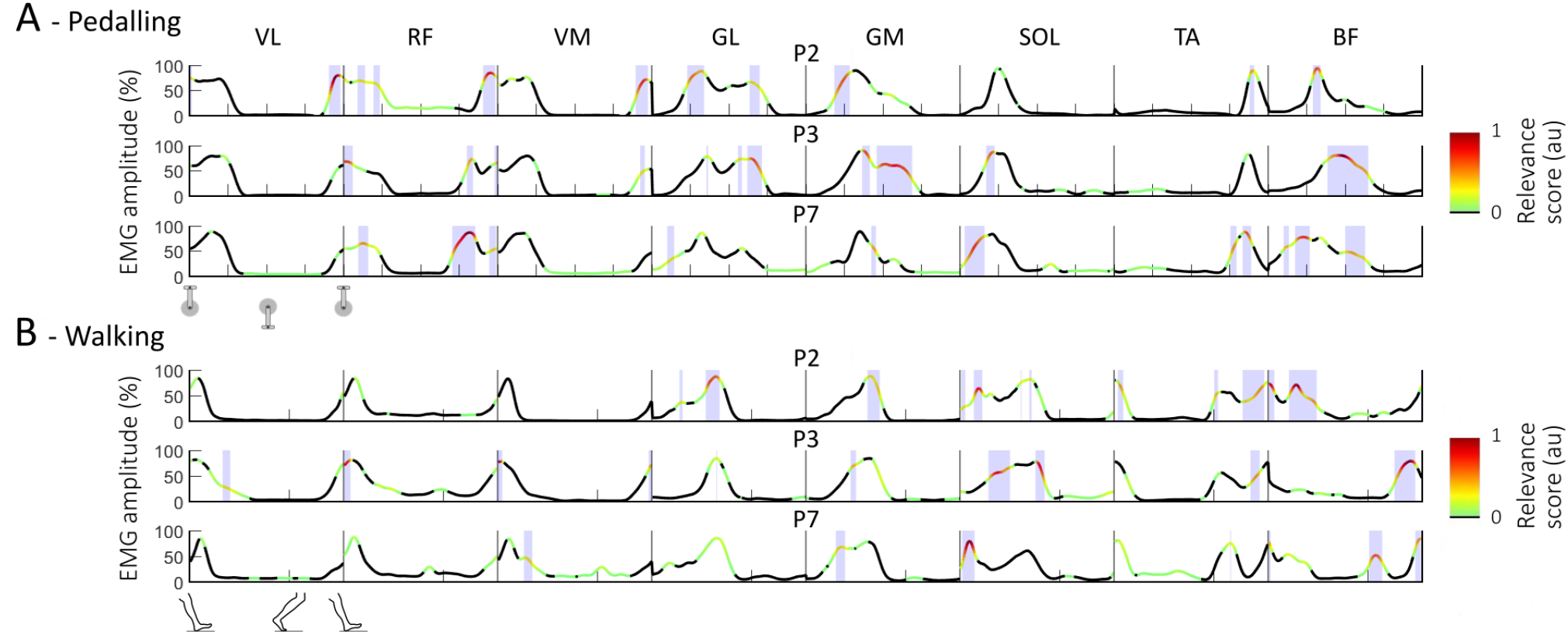
Example results of the relevance scores for three participants (P). The lines show the mean (over 30 cycles) muscle activation patterns for eight lower limb muscles (VL = vastus lateralis; RF = rectus femoris; VM = vastus medialis; GL = gastrocnemius lateralis; GM = gastrocnemius medialis; SOL = soleus; TA = tibialis anterior; BF = biceps femoris) during pedalling (A) and walking (B). The colour bar represents the output relevance scores from the Layer-wise Relevance Propagation technique. Only positive relevance scores, which contribute to the classification of that individual, were considered and therefore all negative and zero scores are coloured in black. The closer the score is to 1, the more relevant the section of the EMG pattern is for recognition of that individual. The shaded areas represent relevance scores greater than the arbitrary threshold of 0.2. The data on the top three panels start with the pedal at top dead centre and the x-axis ticks denote 25, 50, 75% of the pedalling cycle for each muscle. The data on the bottom three panels start at heel-strike and the single x-axis tick denotes the transition between the stance and swing phase.

### Interpretation of Individual Relevance Scores

We first verified if relevance occurred during periods when the muscles were most active. A moderate positive correlation (r = 0.56, p < 0.001) was found between the relevance and relative EMG amplitude. In general, the highest relevance scores were observed during periods of the cycle when the relative EMG amplitude was high. However, high relevance and high relative EMG amplitude were not mutually inclusive; that is, there were many instances of high relative EMG amplitude where relevance was low (Fig. 3). Of note, there were no occurrences of high relevance when the EMG amplitude was low.

**Fig. 3.**
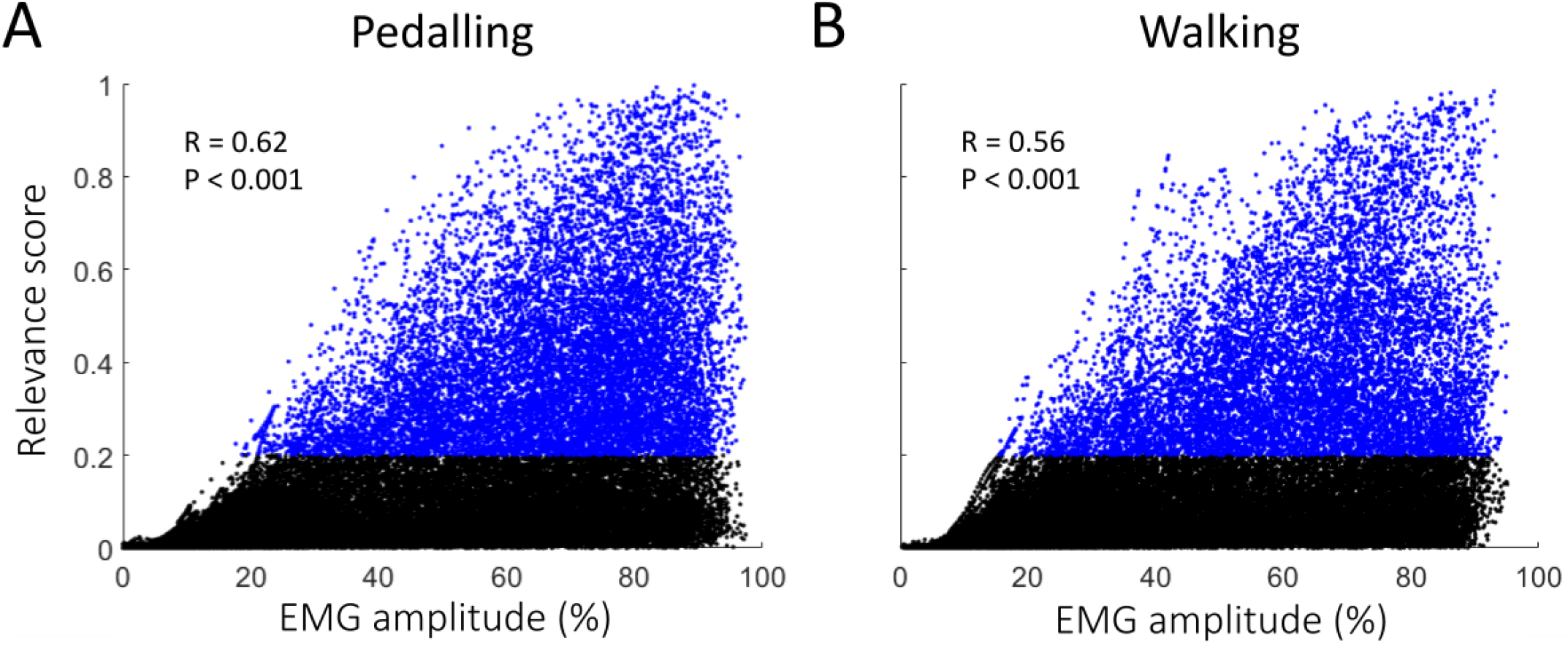
Relationship between EMG and relevance scores. Relationship between the relative EMG amplitude and relevance score for pedalling (A) and walking (B). Each data point from the mean cycle for each participant is shown as a dot. The correlation coefficient and P-value are shown for the pooled data points. Data points for when the relevance scores were greater than the threshold of 0.2 are coloured in blue. The figure highlights that high relevance occurred mostly when the relative EMG amplitude was high. However, there were many instances with high EMG amplitude where relevance was low, meaning that not all phases with high relative EMG amplitude are relevant to participant classification. Of note, there was no high relevance when the EMG amplitude was low. Pearson correlation coefficients were calculated for N = 78. Statistical significance for all correlations was set at *P*< 0.05.

Despite the general pattern of high relevance occurring during periods of high relative EMG amplitude, the periods of the cycle and muscles where high relevance was observed varied greatly between participants. Figure 2 shows the result for three participants. Although all three participants had very similar EMG profiles, there were subtle differences in their timing and shape. These subtle differences were identified by the SVM model during training as being distinctive and unique features that allow classification of the participants. For example, the burst in RF EMG that occurred during the second half of the pedalling cycle was of a similar relative amplitude for all three participants; however, the timing of this burst was slightly different. The results generated by the LRP indicate that the slightly earlier onset observed for participant 7 (Fig. 2A, panel 3) was highly relevant to the classification of this participant and therefore is unique to this participant. In contrast, participant 3 (Fig. 2A, panel 2) seems to have a later and longer activation of both BF and GM during pedalling, which suggests that this pattern is unique to this participant. These results demonstrate the power of the proposed technique, as the models can rapidly extract which features are unique to a given individual (typically <1 min for 78 participants and 30 cycles).

### Robustness of the Relevance Scores

To test the robustness of the relevance scores, we first tested their within-session, between-condition, and between-day reliability for both pedalling and walking. To assess the within-session reliability, we compared the relevance scores across the 30 cycles for each individual. We observed an excellent consistency across cycles, with an average coefficient of correlation> 0.90 (Table 1). However, the root-mean square error (RMSE) values for these comparisons were greater than what would be expected for such high correlation values. This indicates that the magnitude of the relevance scores are less consistent, but that the overall pattern is highly robust. Therefore, we decided to use an arbitrary threshold of 0.2 and considered all magnitudes above this to equally contribute to an individual’s signature.

We further assessed the reliability of the relevance scores across different pedalling conditions (between-condition reliability). To this end, we used the mean relevance per feature (calculated from the 30 cycles) for each participant. The RMSE values were low with very high correlation values (Table 1). Very similar results were also found for the reliability between different days for both pedalling and walking (Table 1).

**Table 1.**
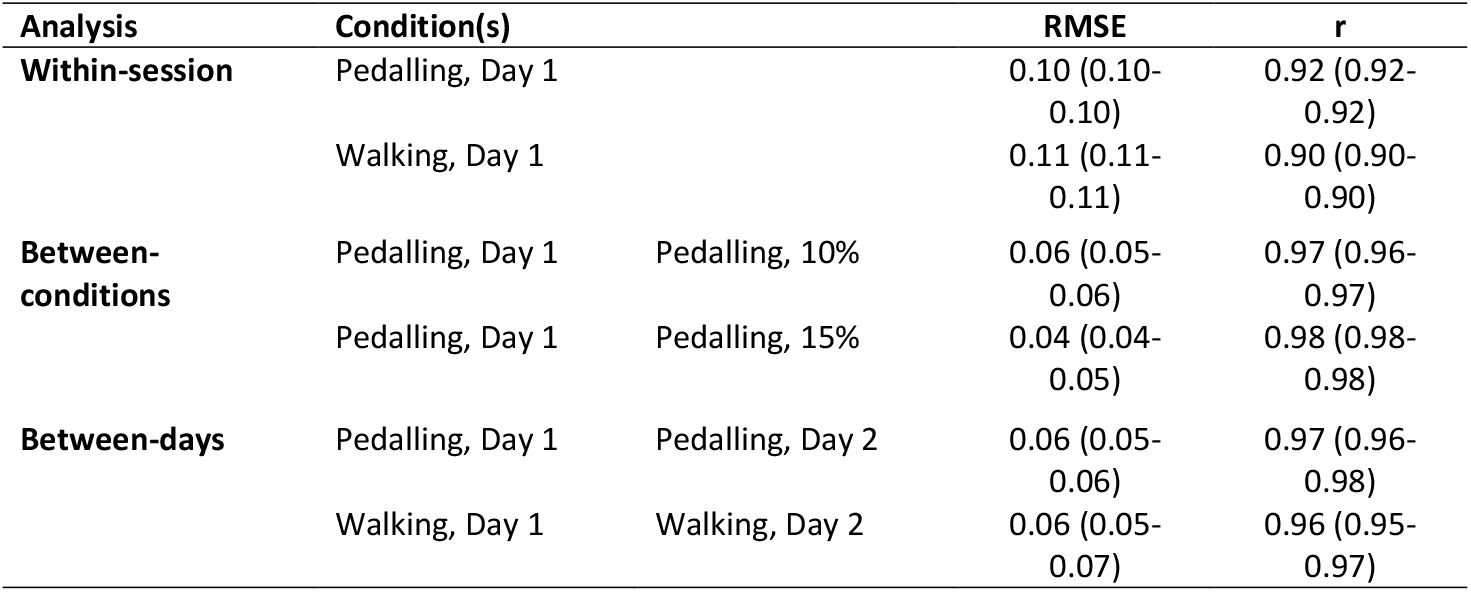
Reliability of the relevance scores. For the between-conditions and between-days, the two conditions per row were compared. Mean root-mean square errors (RMSE) and correlation coefficients with 95% confidence intervals. RMSE numbers are absolute relevance score values, hence they can range between 0-1. Pedalling, Day 1 and Pedalling, Day 2 represent the baseline data from the 150 W condition.

Lastly, we assessed the sensitivity of the relevance scores for the number of participants used for training the models. To this end, we trained several SVM models with the same parameters but with a different number of participants included during training. We used 18 model configurations (i.e. subsample sizes ranging from N = 6 to N = 74 with steps of 4) and compared their relevance scores with the relevance scores from the original model (N = 78 during training, i.e. baseline data; example provided in Fig. 4C). For each of these model configurations, we trained 78 models, one for each participant with N-1 other, randomly chosen participants. We found high correlation values after comparing the mean relevance scores of each participant with the relevance scores from their baseline data (N = 78, Day 1). There was a trend of increasing correlation coefficients with increasing subsample size (ranging from 0.84 for N = 6 to > 0.99 for N = 74 for pedalling (Fig. 4A) and from 0.86 for N = 6 to 1.00 for N = 74 for walking (Fig. 4B)). This indicates that the relevance results are relatively robust even when using a small sample of participants for training of the model. Together with the high consistency found in all the robustness analyses, it suggests that the relevance scores and therefore the muscle activation signatures have a physiological origin rather than being mostly explained by aspects of the methodology.

**Fig. 4.**
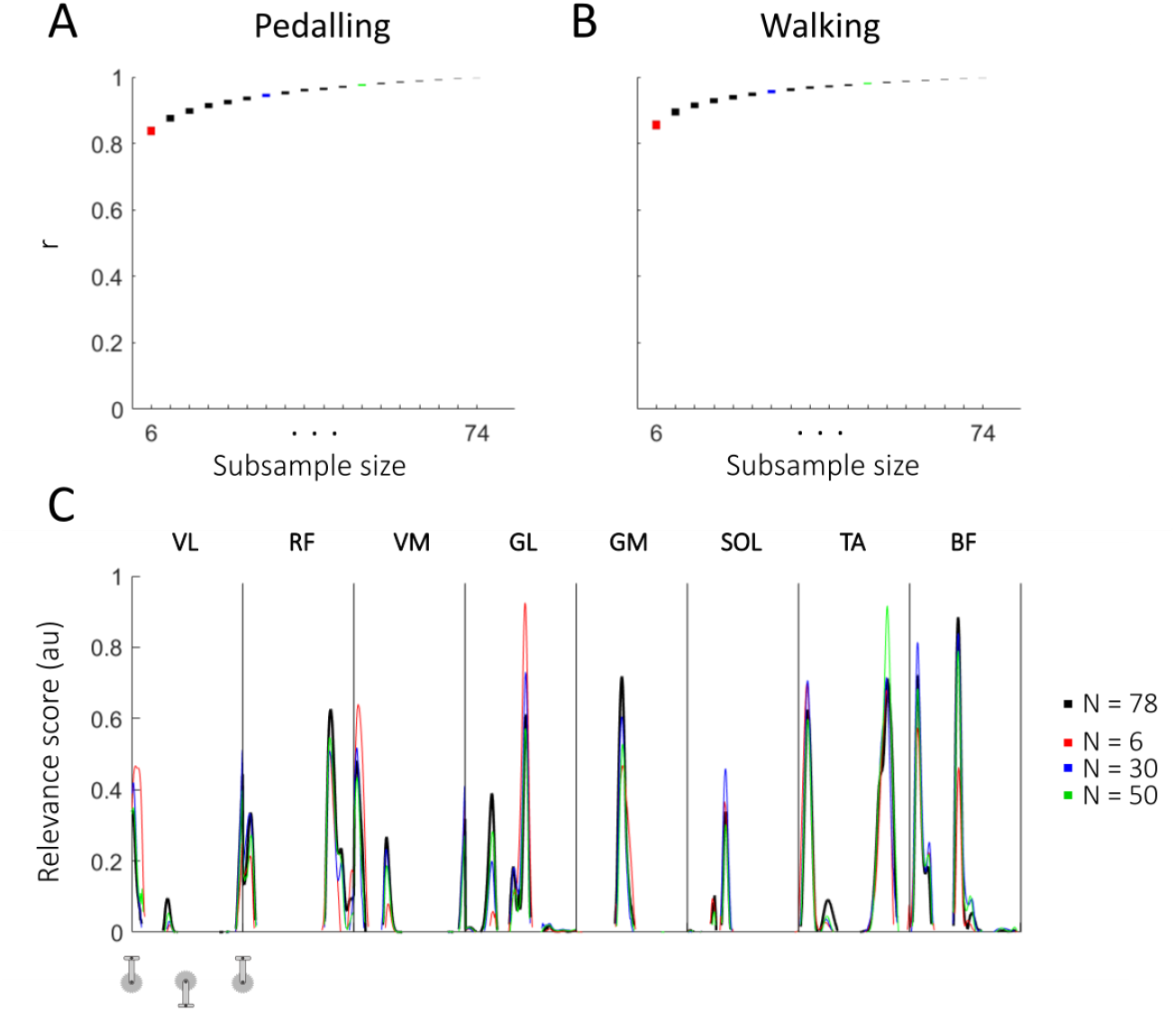
Robustness testing results for different subsample sizes. Correlation results for robustness testing on different subsample sizes using the baseline: Day 1 data for pedalling (A) and walking (B). The mean relevance score extracted for each participant with N = 78 (reference) was compared to the mean relevance score of the same participant extracted from different subsample sizes (for N =6 to N = 74 in steps of 4). Results in panel A and B show the 95% confidence intervals as the y-axis range for each filled rectangle. As an example, panel C shows the relevance scores of a randomly selected iteration and participant. The reference (N = 78) result is shown in black and the results from three different subsample sizes (N = 6 in red; N = 30 in blue; N = 50 in green) are shown. The colours are matched with the subsample sizes of the confidence intervals for the entire group in panel A and B as a reference. Pearson correlation coefficients were calculated for N = 78. Statistical significance for all correlations was set at P < 0.05.

### Determination of Individual Muscle Activation Signatures

Here, we propose a simple methodology for generating a graphical signature map depiction of muscle activation signatures for each individual participant (Fig. 5). For each individual, these signature maps highlight the period(s) of the cycle during which the muscle(s) exhibited a relevance score > 0.2 for all 30 cycles. Signature maps represent a tool that allows for easy and rapid interpretation of each individual’s muscle activation signature, as depicted for all 78 participants in Fig. 6 and Fig. 7). Based on these signature maps, it is clear that the signatures are unique to each participant and task.

**Fig. 5.**
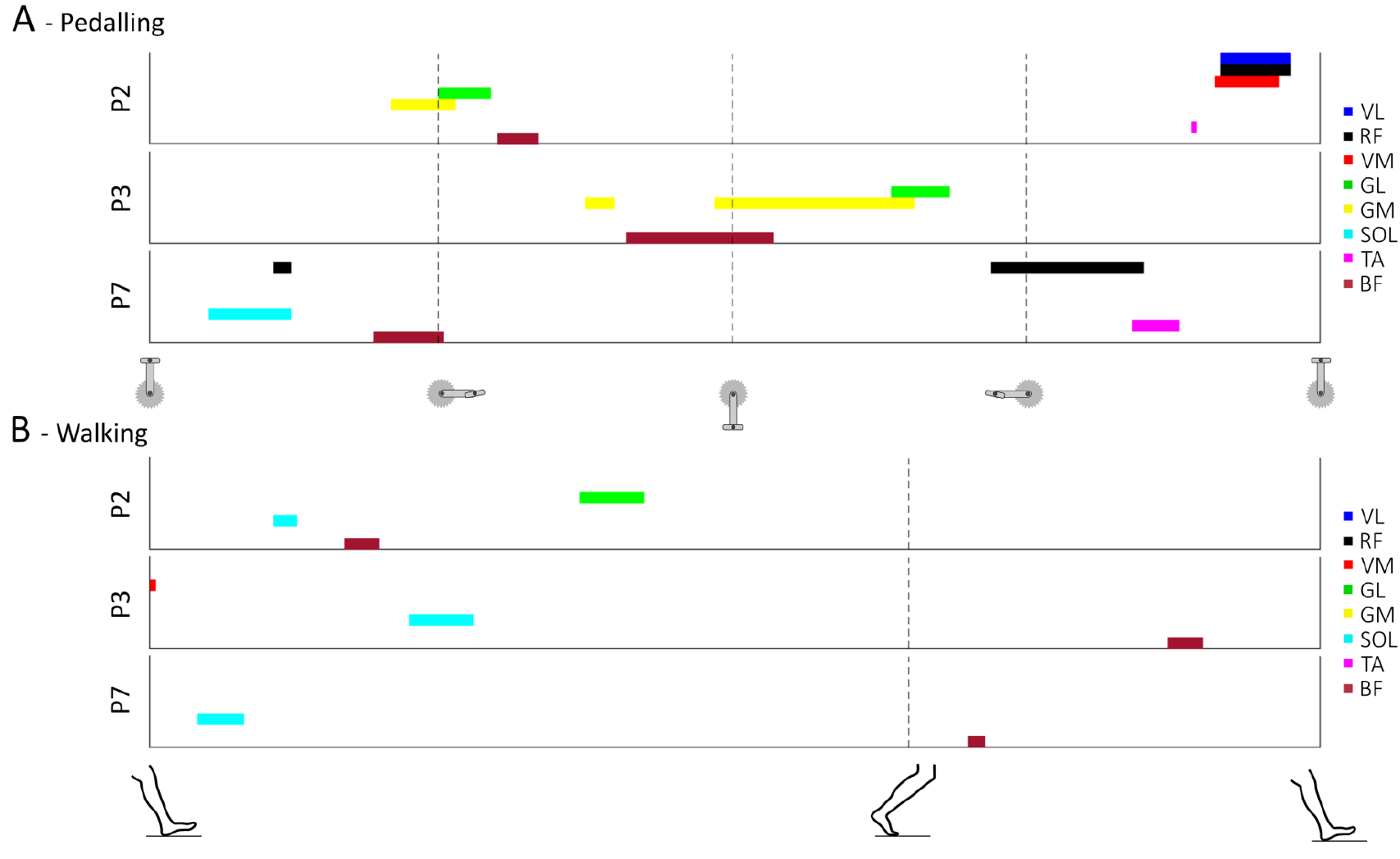
Visualisation of muscle activation signatures. Example muscle activation signature maps for three participants (P) for pedalling (A) and gait (B). Each colour represents a different muscle as shown in the legend on the right. The data on the top three panels start with the pedal at top dead centre and the vertical dashed lines denote 25, 50, 75% of the pedalling cycle. The data on the bottom three panels start at heel-strike and the single vertical dashed line denotes the transition between the stance and swing phase.

**Fig. 6.**
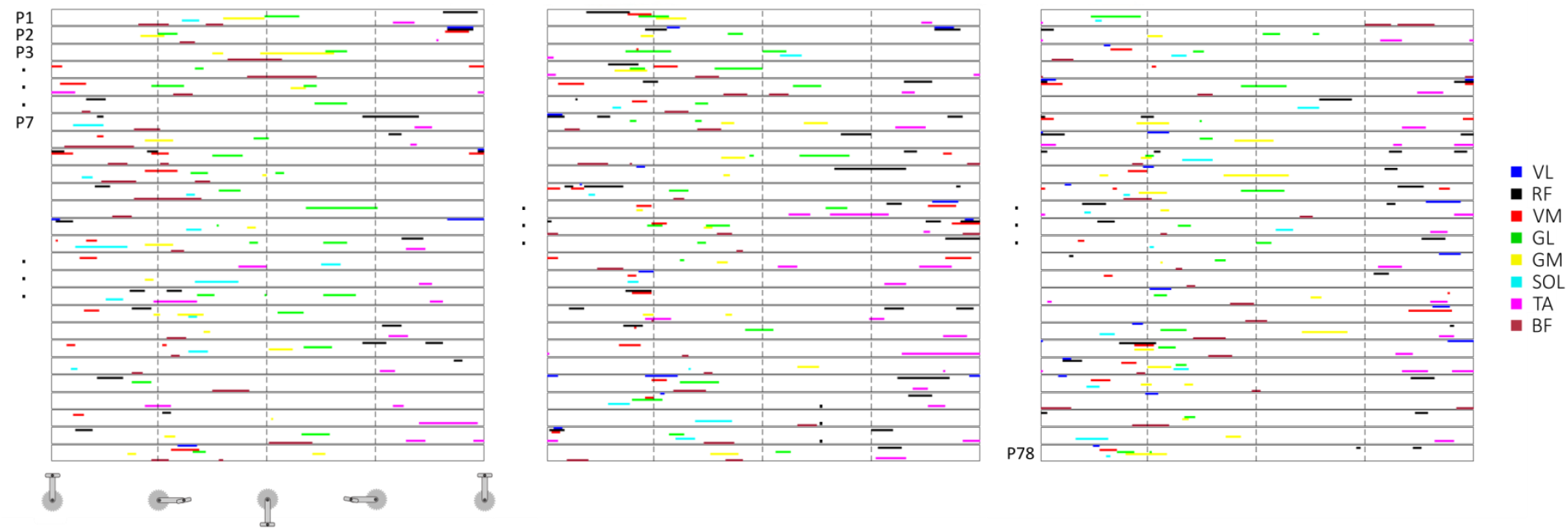
Muscle activation signatures for pedalling. Overview of the muscle activation signature maps of all 78 participants for the baseline pedalling task, Day 1. Each row shows the signature map for a single participant (P), starting with participant number 1 on the top of the left panel and ending with participant number 78 at the bottom of the right panel. The example participants (P2, P3, and P7) for Figures 2 and 4 are indicated. Each colour represents a different muscle, as shown in the legend on the right.

**Fig. 7.**
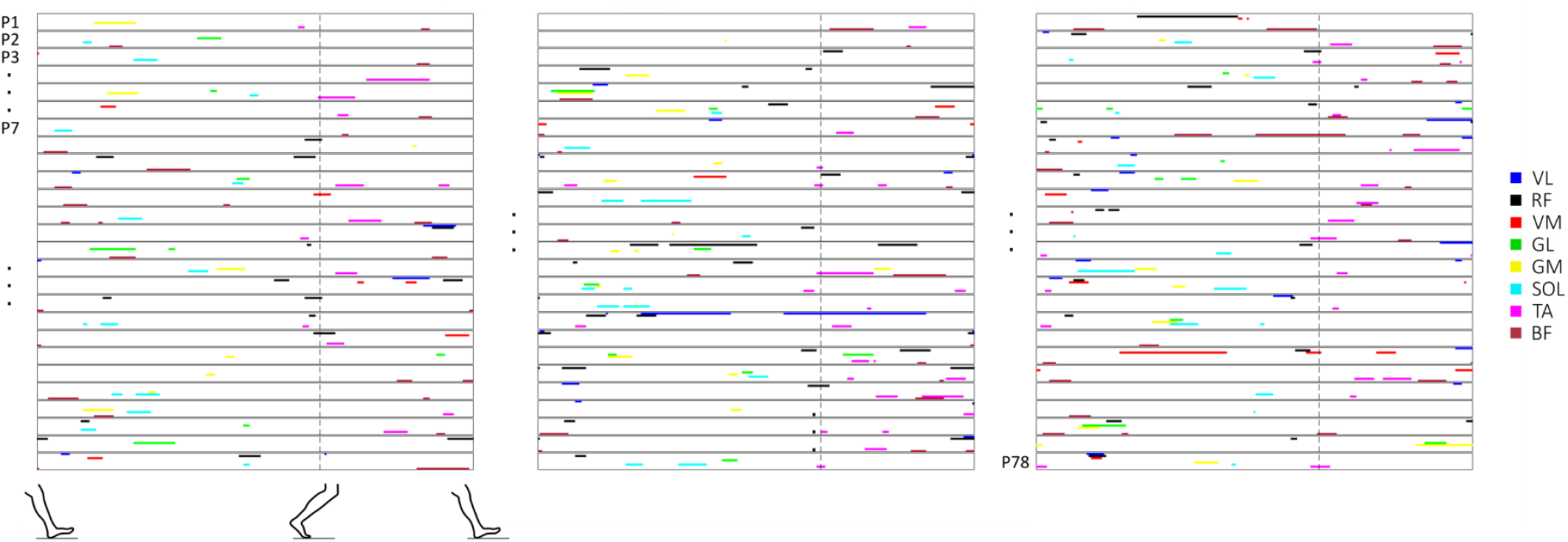
Muscle activation signatures for walking. Overview of the muscle activation signature maps of all 78 participants for the baseline walking task, Day 1. Each row shows the signature map for a single participant (P), starting with participant number 1 on the top on the left panel and ending with participant number 78 at the bottom on the right panel. The example participants (P2, P3, and P7) for Figures 2 and 4 are indicated. Each colour represents a different muscle as shown in the legend on the right.

From these signature maps, we determined which muscles were more often included in the participants’ signatures. This analysis revealed that these muscles differed significantly between participants. Each of the eight muscles was part of the signature of at least several participants (Table 2). However, certain muscles, such as RF, TA, and BF, seemed to be part of the signature of participants more often than other muscles. On the other hand, the GL muscle was often part of the signatures for pedalling but less so for walking, which highlights the notion of task-specificity for an individual’s muscle activation signatures.

**Table 2.**
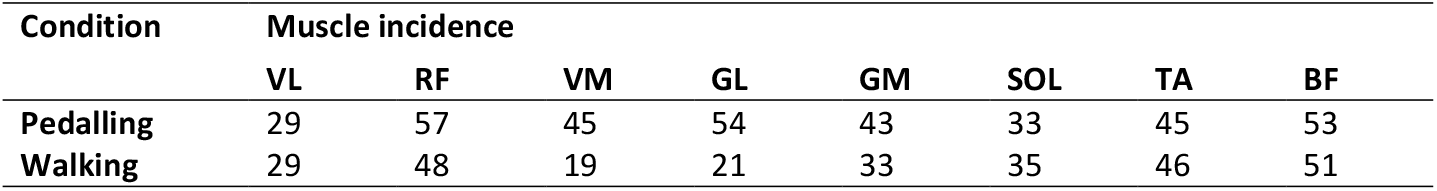
Number of participants (total *N*= 78) for whom each muscle was part of their signature. This analysis was performed using the mean relevance vector per participant for all 78 participants. VL = vastus lateralis; RF = rectus femoris; VM = vastus medialis; GL = gastrocnemius lateralis; GM = gastrocnemius medialis; SOL = soleus; TA = tibialis anterior; BF = biceps femoris.

Fig. 8A shows that the periods during which high relevance scores occurred varied greatly between individuals. High relevance scores for VL and VM muscles occurred mostly around the top-dead centre of the cycle as well as when the pedal was in 90 degrees during the downstroke phase. For walking, we observed that for muscles such as RF, TA, and BF, although the phase of high relevance was spread out over the entire gait cycle, there were two distinct points in time that seemed to contain unique information for many participants, as seen by the darker colours in Fig. 8B.

**Fig. 8.**
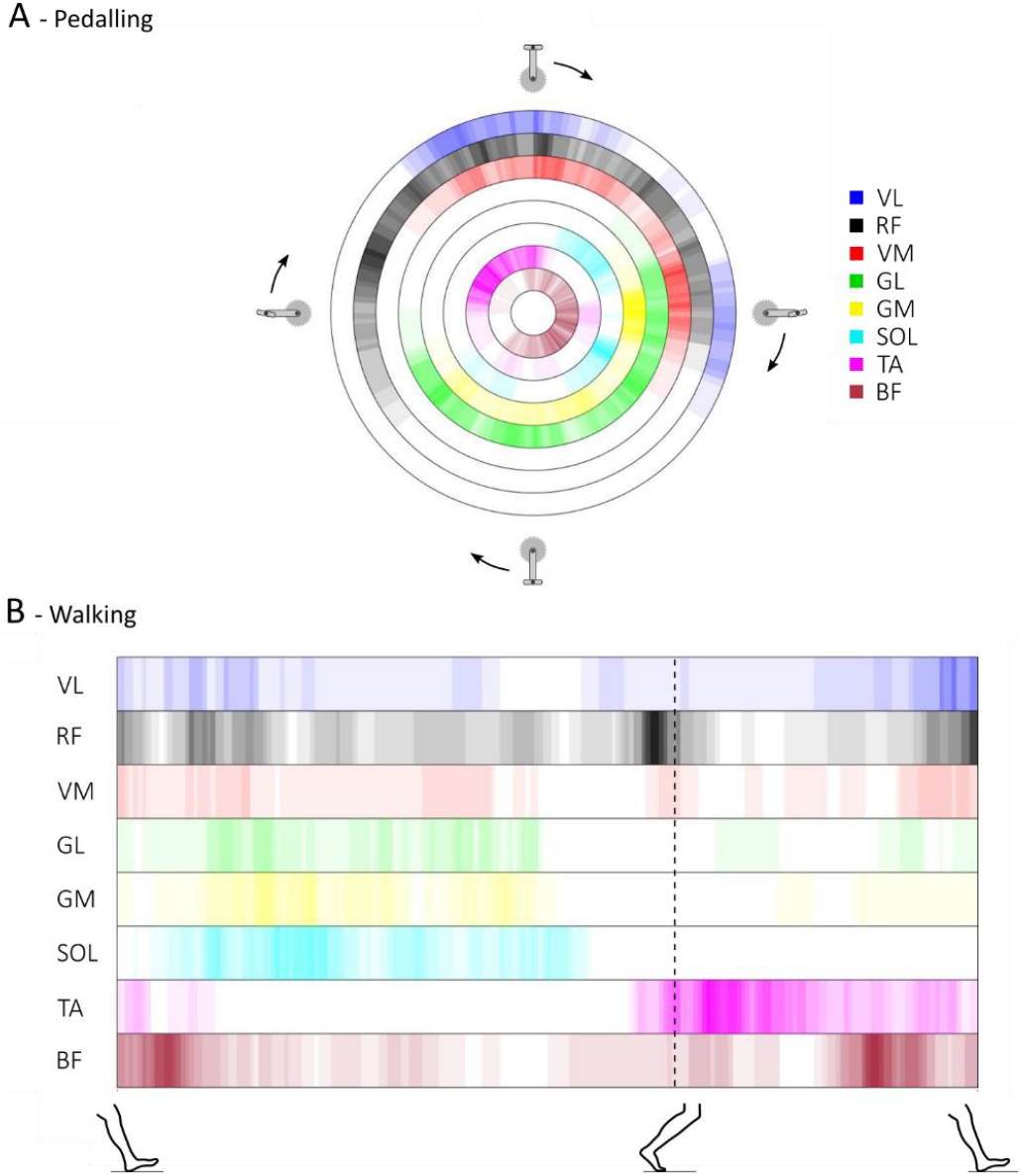
Incidence of features in the signatures. Incidence of muscles in the muscle activation signatures from all participants over the pedalling (A) and walking (B) cycle. Each colour represents a muscle, as shown in the legend on the right side of panel A. The colour gradient represents the incidence of participants having that specific muscle as part of their signature at that point of the cycle, with lighter colours representing less participants and darker colours representing more participants. The maximal incidence (darkest colour; i.e. the colour used in the muscle legend) is set at N = 15. The figure provides a graphical overview of all the signatures from the testing group (baseline, Day 1; N = 78) and highlights which phase of the cycles the muscles were part of the signatures of the tested participants. Areas coloured white indicate that no single participant had the specific muscle as part of their signature during that specific phase of the cycle. For pedalling, the figure shows that the timing of the signature features relative to the pedalling cycle varied significantly between muscles. For example, some muscles, such as VM and VL, were part of the signature only between the end of the upstroke and just past halfway through the downstroke, with the majority of incidence occurring during the first and final sections of this phase, as seen by the darker colours. Other muscles, such as RF or BF, were part of the signatures for a longer section of the cycle, starting halfway through the first part of the upstroke until the second part of the downstroke. This indicates that the variation in the phase of the cycle during which unique features occurred for this data set varied more RF and BF. For gait, the data is more spread-out over all muscles except for SOL and TA, with no participants having unique features in their signature during the swing phase for SOL and no participants for TA during roughly 25-75% of the stance phase.

## Discussion

Although variability of muscle activation strategies between different individuals is well-documented, the idea that each individual exhibits unique activation strategies is relatively novel. Based on the observation that machine learning algorithms can accurately identify individuals based on their muscle activation patterns, Hug et al. (9) suggested the existence of individual muscle activation signatures. However, Hug et al. (9) did not provide information regarding the decisions made by the algorithms in making a particular classification, making it impossible to provide conclusive evidence of the physiological origin of the signatures. In the current work, we developed an approach based on the use of LRP that allowed us to map out individual muscle activation signature of 78 participants. The robustness and low sensitivity to the trained model (i.e. the low influence of the combination and number of participants used to train the model) of our results provide strong evidence of a physiological origin of the activation signatures and thus of their physiological and potentially clinical relevance. Identification of these signatures is a crucial step toward a better understanding of human movement in the context of health and disease.

### From Relevance Scores to Muscle Activation Signatures: methodological considerations

The LRP technique applied to interference EMG signals demonstrated which muscles and associated time periods were the most relevant for identifying an individual during cycling or walking by providing time-varying relevance scores. Verifying the reliability of these relevance scores was a prerequisite to confirm the existence of physiological signatures of movement production (Table 1). First, we confirmed the consistency of the relevance scores between cycles when tested on the same day, during the same task/condition, and with the same electrode placement. Second, we demonstrated the consistency of the relevance scores when they were extracted from cycling at different intensities. Third, to rule out the possible effect of electrode placement, we tested the robustness of the signatures between two separate days. We observed that the relevance scores had high reliability. Finally, we demonstrated that the relevance scores were relatively insensitive to the number of participants used to train the models (Fig. 4). Together, these analyses provide strong evidence that the signatures originate from physiological features unique to each individual rather than from methodological features.

In this study, the muscle activation signatures were extracted from interference EMG. In order to understand the physiological meaning of the features extracted from these signatures (derived from the EMG patterns of eight muscles), it is important to consider the interpretation of interference EMG amplitude as a neural strategy. First, it is well-known that EMG signals are affected by non-physiological factors such as the thickness of the subcutaneous fat layers, the shape of the volume conductor, and the location of the electrodes (14). To minimise the impact of these non-physiological factors, we normalised the EMG signals to the peak EMG amplitude measured during the task (15). In other words, we considered the time-varying profile of the EMG signals without any information regarding their actual magnitude with respect to maximal activation. Second, it is important to note that interference EMG does not provide direct information about the neural drive; i.e. the number of motor neuron action potentials (16, 17). Instead, interference EMG amplitude is more closely related to muscle activation, which is related to the number of muscle fibre action potentials (16, 17). Note that the relationship between neural drive and muscle activation depends on the size of the active motor units; i.e. the number of muscle fibres within each motor unit. Based on these considerations, we considered the normalised time-varying EMG profiles to be time-varying muscle activation profiles and interpreted our results as the existence of individual muscle activation signatures. Future work is needed to confirm that these signatures originate from unique neural strategies.

### Neuromechanical Coupling to Explain the Uniqueness of Muscle Activation Strategies

There is strong coupling between the neural drive to a muscle and several biomechanical factors that influence the muscle’s force generation (18, 19). In other words, muscle activation may be tuned to each individual’s specific anatomy and to the biomechanical properties of their musculoskeletal system. For example, tendon compliance affects the time required for force transmission between the onset of muscle fibre shortening and torque production. It varies substantially (up to 60%) between individuals and between different muscle-tendon units (20). To account for the longer delay in force transmission in a more compliant tendon, the muscle would have to be activated earlier or more (or both) as compared to a muscle with lower in-series tendon compliance. Furthermore, the different combinations of tendon compliance and muscle architecture generally observed between individuals (21) likely further induce variation in the timing and magnitude of muscle activation (22). Similarly, an individual’s anthropometry will affect segmental inertial properties and muscle moment arms involved in joint rotations, which in turn may affect the requirements of muscle activation (23). Such anatomical and biomechanical variations between individuals would give rise to unique differences in their time-varying patterns of activation in order to meet the requirements of force production for motor tasks. Of note, it is possible that the influence of these structural and biomechanical features on the time-varying EMG profiles was partly cancelled out by our normalization procedure. However, this normalization procedure affected only the amplitude of activation, keeping the interindividual differences in activation timing unchanged

It was recently postulated that motor unit recruitment, at least in respiratory muscles, may not follow the generally-accepted size principle in which motor units are recruited in order of increasing size (24), but rather that they are recruited with respect to their mechanical advantage (25). Given the high variability in muscle biomechanical properties between individuals, it is likely that muscle activation patterns are also tuned to account for these degrees of freedom. The constant nature of these factors over the short term would explain why these unique activation signatures are robust within each individual. It is important to note that a causal relationship between mechanical and neural aspects of muscle contraction has not been established and it remains possible, although less likely, that variability of muscle properties among individuals is due to different activation strategies. Regardless of the cause-effect order, there is a large amount of evidence for neuromechanical coupling, which provides a plausible explanation for the observed muscle activation signatures.

### Biomechanical and Neurophysiological Interpretations

High relevance scores occurred during periods of high relative EMG amplitude, but not all periods of high relative EMG amplitude were relevant for participant classification (Fig. 3). Indeed, the classification model considered only a few phases of the cycle from only a few muscles (Fig. 5 and 6). Of note, the periods of high relevance differed substantially between pedalling and walking. For example, the VM and GL muscles were far less frequently part of the signatures in walking compared to pedalling (Table 2). Fig. 5 further highlights these differences in a few participants. Given the substantial differences in both kinematics and kinetics between walking and pedalling, such differences are to be expected. Even though the muscles used for classification varied greatly between tasks and between participants, three muscles (RF, BF, and TA) were used more than the other muscles for both pedalling and walking. This observation is consistent with our current knowledge of the biomechanical role of these muscles, since the bi-articular RF and BF muscles mainly transfer energy between joints. Together with the TA muscle, these muscles contribute to the stabilization or control of the motion, which is considered to be a secondary role as opposed to contributing to power production (26). This is consistent with the observation that the degree to which a motor component varies between individuals depends on the role of that motor component in movement (27), and that the variability in neural control between participants is high when the effect on motor output is low (28).

Several studies have proposed that individuals have their own “motor program styles”, with considerable variation between individuals in motor control strategies, including muscle synergies, but with high consistency within each individual (29). Such variation in motor control strategies has been shown for movement in both humans (30) and other animals (31) and is consistent with our findings of robust individual muscle activation signatures. Large variation in neural responses between individuals has also been reported for odour discrimination tasks in rats (32) and in the reotaxic response of sea lamprey (Petromyzon marinus L.; 33). However, it is important to keep in mind that the existence of such an inter-individual variability does not prove the existence of individual signatures. Using inter-individual differences as evidence of personal signatures requires two important considerations: First, these strategies should be robust across multiple sessions. Second, these strategies should not be exactly the same for any two individuals, i.e. they should be unique. In the current study, we ensured that the activation strategies were robust and unique. Therefore, to the best of our knowledge, the current study is the first to provide evidence that individuals use unique activation (and likely neural) signatures. Collectively, these findings suggest that unique features may drive the neural strategy for each individual in several biological processes in both animals and humans. This emphasises the need to carefully study individual patterns rather than losing useful information by averaging data from a group of individuals. This can provide important knowledge that will facilitate a better understanding of the fundamental principles behind the neural control of movement.

### Applications and Future Directions

One strength of our approach is that it can detect small differences in EMG patterns between individuals that would likely be missed by a human observer. It is important to note that it is the combination of the high relevance periods of all eight muscles that resulted in highly accurate classification, and thus constitutes an individual’s muscle activation signature. Finding such complex patterns with these many degrees of freedom is highly difficult for a human observer, as it would require keen perception combined with expert domain knowledge and a detailed understanding of the data. In this work, we proposed a simplified depiction of individual muscle activation signatures (Fig. 6), which allows a human observer to make fast and easy comparisons between individuals, with possible clinical applications. Surface EMG is widely used in research settings and in clinical routines to identify abnormal activation patterns. Although current automated approaches can distinguish between healthy and pathological EMG signals (34), these approaches cannot be used for differential diagnoses, especially in complex neural disorders. This is mostly because of the large variability found between individuals and our relatively poor understanding of the relevance of this variability. Our approach allows for this inter-individual variability to be investigated more in depth. Future research should therefore aim to implement the proposed approach for pathological populations and to verify its validity and effectiveness in clinical settings.

Multiple scientific domains are transitioning toward personalised approaches where diagnosis or treatment are based on individual characteristics (35). Such personalized approaches are crucial to improve our understanding of movement in healthy populations (36) as well as in populations with neurological disorders (37). Importantly, these approaches require the identification of an individual’s unique features, but this is rarely performed for muscle activation. Together with subject-specific musculoskeletal models, the unique activation features could be used to explore the effect of highly-specific and realistic combinations of neural and biomechanical parameters on movement. This is an important future step, since the results from this study cannot be used to interpret whether these features have any direct influence on movement kinematics or kinetics. Due to the complex interplay between neural and biomechanical features during movement (38, 39), taking this step will represent a major advancement in our ability to understand, prevent, and treat such neurological disorders. The proposed technique also has profound implications in the optimisation of bio-inspired exoskeletons by optimising these devices to individual muscle activation strategies (40). Another fast-evolving area that could indirectly benefit from our approach is the artificial control of weakened or paralysed muscles through electrical stimulation (41). In order to provide optimal electrical stimulations, we need to first better understand the biological importance of unique individual muscle activation patterns. Thus, there are plenty of opportunities for the suggested technique to contribute to the significant advancement of current scientific practices.

## Funding

François Hug is supported by a fellowship from the *Institut Universitaire de France* (IUF). Sebastian Lapuschkin is supported by the German Ministry for Education and Research as BIFOLD (refs. 01IS18025A and 01IS18037A) and TraMeExCo (ref. 01IS18056A). Support was received from the French national research agency (ANR-19-CE17-002-01, COMMODE project; to FH).

## Author contributions

JA performed the analyses and interpreted the results, wrote the manuscript, created the figures, and approved the final version of the manuscript. FaH performed the analyses and interpreted the results, wrote the manuscript, and approved the final version of the manuscript. SL performed the analyses and interpreted the results, wrote the manuscript, and approved the final version of the manuscript. LL conceived the experiment, collected the data, and approved the final version of the manuscript. FH conceived the experiment, collected the data, performed the analyses and interpreted the results, wrote the manuscript, and approved the final version of the manuscript.

## Competing interests

The authors have no competing interests in this work, financial or otherwise.

## Data and materials availability

All raw EMG data are available as supplemental material at https://doi.org/10.6084/m9.figshare.8273774. Python and Matlab scripts are available at https://github.com/sebastian-lapuschkin/interpretable-emg-signatures.

## Notes

https://doi.org/10.6084/m9.figshare.8273774

### Competing Interest Statement

The authors have declared no competing interest.

### Summary of Updates

This is the final version that was submitted after revision to Journal of The Royal Society Interface.

